# How to Become Invisible to Mosquitoes: A Computational Study of Host Signal Collapse

**DOI:** 10.64898/2025.12.09.693171

**Authors:** Zi-Ru Jiang

## Abstract

Mosquitoes locate vertebrate hosts by following coherent gradients of carbon dioxide, heat, and humidity. Nevertheless, a persistent empirical observation is that ordinary electric fans drastically reduce mosquito bites, even in the absence of chemical repellents. Here, we introduce a physics-guided, agent-based simulation demonstrating that mosquito host-seeking is not chemically inevitable but physically fragile. The model integrates directional airflow, vortex-induced chaotic mixing, and thermal decoy fields, and tracks 250 autonomous mosquito agents over 600 time-steps under Monte-Carlo sampling with independent randomized initializations. Three aerodynamic mechanisms were quantified. Mild upward airflow already reduces successful host localization by more than 80%, despite not mechanically preventing flight. Increasing vortex circulation produces a continuous, threshold-like collapse, reducing localization probability below 1% at moderate strength. Thermal decoys cause only a linear dilution of success, indicating misinformation alone cannot trigger collapse. A two-dimensional phase map reveals a robust “invisibility region” where airflow and vortex perturbations interact synergistically, eliminating host detection even when neither factor individually reaches threshold. These results show that mosquito host-seeking operates as a gradient-dependent failure system: once scalar fields lose spatial coherence, navigation collapses independent of sensory capability. This provides a quantitative theoretical basis for fan-mediated mosquito protection and suggests that non-chemical, low-energy airflow strategies can induce physical invisibility to hematophagous insects.

## 1. INTRODUCTION

Mosquitoes remain the world’s deadliest animals, responsible for more than 700,000 human deaths each year through malaria, dengue, chikungunya and other vector-borne diseases (World Health Organization, 2023). Although long-lasting insecticidal nets (LLINs), indoor residual spraying (IRS), and emerging gene-drive or Wolbachia-based approaches have significantly reduced transmission, global malaria burden has plateaued since 2015 and is now resurging across high-transmission regions (Bhatt et al., 2015; Lawal et al., 2024; Weiss et al., 2025; Umugwaneza et al., 2025). Widespread pyrethroid resistance, declining insecticide durability, and escalating operational costs increasingly indicate that chemical or genetic interventions alone are insufficient for achieving long-term elimination (Ranson & Lissenden, 2016).

In parallel, there is growing interest in low-cost, non-chemical environmental manipulation, particularly airflow. Field studies have shown that ordinary household fans can substantially reduce mosquito landings and biting indoors, even without repellents, apparently by disrupting host-seeking behavior (Sutcliffe & Yin, 2021; Carrasco-Tenezaca et al., 2023). However, airflow remains absent from global vector-control frameworks, and major reviews still treat it as a laboratory nuisance rather than a behaviorally meaningful suppressor (Kline, 2006). This persists despite convergent experimental evidence demonstrating that airflow significantly modifies mosquito navigation, sensory integration, and plume detection.

Mechanistically, airflow impacts host seeking on several sensory and physical levels. Wind-tunnel experiments show that airflow distorts CO2 plume topology, filament intermittency, and gradient stability, thereby impairing chemotactic tracking and heat localization (Cooperband & Cardé 2006; McMeniman et al., 2014; Sutcliffe & Yin, 2021; Carrasco-Tenezaca et al., 2023; Hinze et al., 2021). Complementing these findings, Cribellier et al. (2018) demonstrated that even in the absence of bulk wind, local recirculation vortices around odour-baited traps fragment odor filaments, generate intermittent “lost–found–lost” trajectories, and induce repeated reorientation events—direct behavioral evidence that small-scale turbulence alone can degrade plume coherence and destabilize host-approach paths. Beyond olfaction, recent work demonstrates that thermal and olfactory cues act jointly, with human scent strongly shaping mosquito thermotaxis via conserved sensory pathways (Giraldo et al., 2023). These findings integrate with broader multisensory reviews indicating that mosquitoes use a rich suite of olfactory, thermal, hygrosensory, visual, and inertial cues to navigate complex environments (Montell, 2025). Even background CO2 concentration can significantly alter search performance, pointing to a critical role for plume geometry, not just odor intensity (Majeed et al., 2014).

Despite these advances, airflow has typically been treated experimentally as binary “low vs. high” perturbations, without quantifying why host detection sometimes declines gradually whereas in other cases it collapses abruptly. Three fundamental questions therefore remain unresolved: What determines the collapse threshold at which host detection probability falls below 1%? Do directional airflow and vortex turbulence suppress host seeking through distinct mechanisms? Do airflow and thermal misinformation combine additively or multiplicatively?

Recent developments in plume physics and search theory conceptualize mosquito host seeking as an information-limited gradient-climbing process, rather than a deterministic olfactory response. Similar to bacterial chemotaxis and theoretical infotaxis (Vergassola et al., 2007), mosquitoes track scalar fields whose structure is highly vulnerable to turbulence. Stretch–fold– shear cascades can destroy scalar coherence faster than insects can integrate temporal cues (Shraiman & Siggia, 2000). Under this framework, airflow may render a host functionally invisible despite mosquitoes retaining full flight ability and intact sensory systems—a hypothesis directly supported by showing sharp behavioral redistribution and strong suppression of host-contact behavior at low wind speeds (Sutcliffe & Yin, 2021).

Although prior studies have independently examined directionality (Hoffmann & Miller, 2002), ventilation-driven suppression (Carrasco-Tenezaca et al., 2023), or turbulence-induced fragmentation (Lei et al., 2023), no unified mechanistic simulation has yet integrated directional airflow, vortex mixing, and thermal misinformation under a common physics-based ruleset. As a result, the field lacks an integrative explanation for the nonlinear—sometimes catastrophic— failure of mosquito host seeking in disturbed airflow environments.

To address this gap, we developed a physics-guided, agent-based simulation integrating: (i) 2D CO2 and heat scalar fields governed by a 1/(1+r) decay model; (ii) 250 autonomous mosquito agents performing noisy gradient-biased walks; (iii) three aerodynamic perturbation modules— uniform drift, vortex-induced mixing, and thermal-decoy fields; (iv) 30–50 Monte-Carlo runs (50 used unless otherwise stated) repetitions per parameter set (>2.4 million total flights). This unified framework enables quantification of collapse thresholds, comparison of aerodynamic suppression modes, and identification of how turbulence, directionality, and thermal misinformation interact to determine whether a host remains detectable—or becomes aerodynamically invisible.

## 2. METHODS

### 2.1 Model Overview

We developed an agent-based navigation model to quantify how airflow and vortex-like perturbations disrupt odor–heat guided host-seeking flight. The model simulates *N* mosquitoes moving within a 2D arena toward a stationary host that emits a chemothermal gradient. At each discrete time step, individuals update their heading based on the combined influence of (i) deterministic CO2–heat gradients, (ii) deterministic aerodynamic forcing (wind or vortex), and (iii) stochastic sensory noise. To focus solely on navigational information, the model does not include mechanical blow-off or loss of flight stability; all mosquitoes remain fully flight-capable under all airflow intensities, consistent with classical observations that mosquitoes can sustain upwind flight at >1–1.5 m s^−1^ when odor cues remain intact (Kennedy, 1940; Gillies & Wilkes, 1970). The main output variable is the proportion of mosquitoes that reach the host region within a fixed time horizon. All simulations are performed using the Python implementation in *mos_phase_models_v20*.*py*. This architecture isolates deterministic environmental structure from stochastic sensory uncertainty, enabling clear identification of airflow-induced collapses.

### 2.2 Spatial Domain and Host Field Construction

A square lattice of size GRID × GRID (default 256×256) defines the navigable space. The host is positioned at the center (HOST = [GRID/2, GRID/2]). Two scalar fields are precomputed following the script’s build_fields () function:

Thermal field:

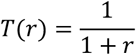

CO2-like odor field:

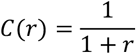

Combined attractant:

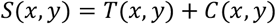

Centered finite differences:

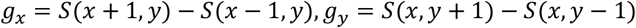

implemented as:

gx = total[x+1, y] - total[x-1, y]

gy = total[x, y+1] - total[x, y-1]

grad_move = np.stack([gx, gy], axis=1)

The gradient term provides the deterministic chemothermal drive.

### 2.3 Deterministic Aerodynamic Forcing

Two airflow regimes were examined independently.

#### (a) Uniform Wind Field

A spatially uniform advective field from calc_wind_velocity():

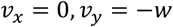

#### (b) Vortex Field

Rotational airflow from calc_vortex_velocity():

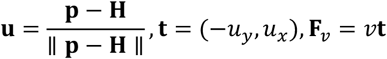

The vortex field represents a simplified, deterministic analogue of the recirculation zones and odor-filament fragmentation documented around odour-baited traps (Cribellier et al., 2018), where small-scale rotational flow causes intermittent plume contact and repeated orientation failures. Both airflow components are strictly deterministic; no random gusts or turbulence realizations are included, allowing threshold behavior to be attributed solely to aerodynamic structure.

### 2.4 Sensory Noise

Sensory noise is modeled as additive Gaussian perturbation applied to the movement vector: noise_move = SENSORY_NOISE * rng.normal(size=grad_move.shape)

This is the only source of stochasticity; no randomness exists in the airflow or the scalar fields.

### 2.5 Movement Update Rule

At each time step:

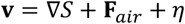

Movement is direction-dominant:

move = grad_move + flow_move + noise_move

step = np.sign(move)

pos = pos + step

Positions are clipped within the arena. Airflow alters heading direction only; it does not increase step magnitude, ensuring that collapse reflects loss of navigational information rather than forced kinematics. This directional update emphasizes heading errors rather than speed modulation, representing a worst-case navigation regime.

### 2.6 Initialization and Success Criterion

At the start of each simulation, N_MOSQ_ = 250 mosquito agents are initialized at random positions within the 2D arena:

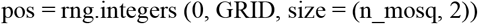

Each simulation proceeds for a maximum of 600 discrete time-steps, unless all agents have either reached the host or exited the effective search region. An individual mosquito is classified as successful if, at any time during the simulation, its position enters a circular target zone of radius 8 pixels centered on the host location.

The success rate for a single simulation is defined as:

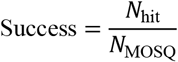

where *N*_hit_ denotes the number of agents that reached the host zone.

### 2.7 Monte-Carlo Protocol and Phase Scans

To characterize the effect of airflow parameters, we conducted Monte-Carlo simulations across a predefined parameter space. For each wind or vortex intensity level in the interval [0, 3] with increments of 0.1, we performed 50 independent realizations, each using a distinct but reproducible random seed to ensure statistical robustness. Unless otherwise specified (e.g. Fig. S1), all main-text figures use 50 Monte-Carlo runs.

For a given airflow parameter *w* (wind) or *v* (vortex), the mean success probability is computed as:

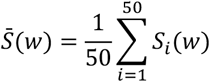

where *S*_*i*_(*w*) is the success rate observed in the *i*-th simulation. The ensemble-averaged values 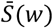 and 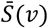 were used to construct the phase diagrams presented in the main text.

### 2.8 Rationale for Model Structure

The model isolates airflow–gradient interactions while minimizing biological and computational complexity. Key simplifying assumptions (kept minimal): 2D navigation (indoor CO2 plumes are quasi-planar); Static scalar fields; no temporal fluctuations; No inter-agent interactions; No additional sensory modalities (visual, humidity, skin volatiles); No mechanical blow-off, tumbling, or flight loss.

These assumptions ensure that observed failures originate from physical destruction or displacement of the chemothermal gradient, not from extraneous behavioral mechanisms. This abstraction is justified by empirical evidence showing that turbulence-induced intermittency— not mechanical impairment—is the primary driver of navigation failure near vortex-dominated flows (Cribellier et al., 2018; Sutcliffe & Yin, 2021). Airflow effects are implemented as distortions of agents’ gradient sampling rather than explicit advection of the scalar fields, isolating information loss from full plume dynamics. Here, “entropy dilution” refers to the flattening of the scalar attractor landscape without removal of its global maximum.

## 3. RESULTS

### 3.1 Weak wind perturbation triggers an abrupt navigational collapse (without crossing the invisibility threshold)

Across all tested wind strengths, host-seeking success exhibited a highly nonlinear, threshold-like drop (Fig. 1). A minimal airflow of w=0.3 caused an 82% reduction (0.34→0.06), and success reached a low plateau near ~0.045 above w≈1.0. The fact that near-collapse occurs at very weak airflow agrees with wind-tunnel experiments showing that *Aedes aegypti* approach rates can fall by 60–70% at only 0.2–0.3 m/s (Sutcliffe & Yin, 2021). Although wind alone does not cross the invisibility threshold (*P* < 0.01), it induces a near-collapse plateau that renders host detection functionally unreliable.

**Figure 1.**
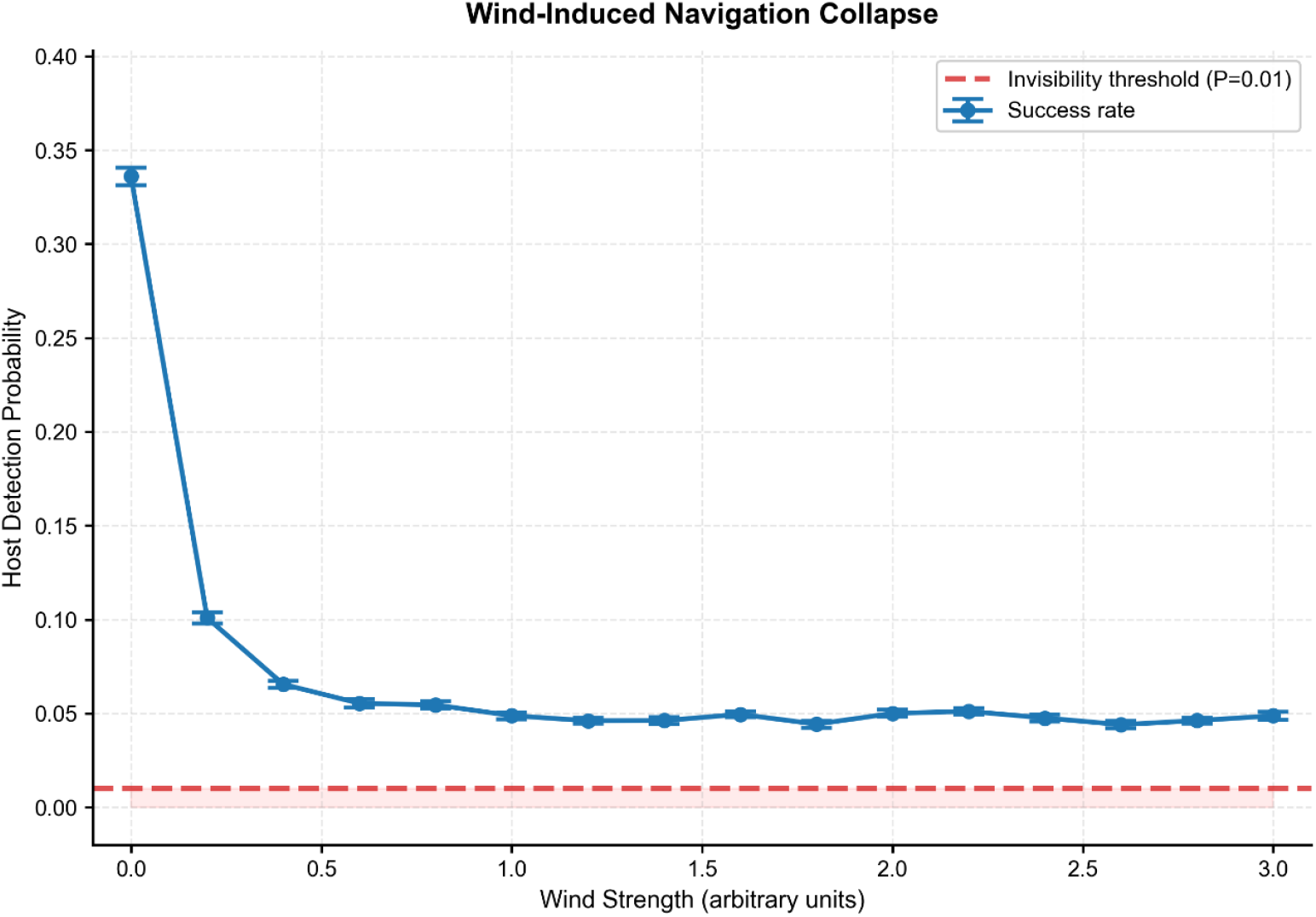
Effect of wind strength on mosquito navigation success. Mean host-detection probability declines sharply from 0.34 at zero airflow to 0.06 at wind strength w=0.3, representing an 82% reduction in successful host localization. Beyond w>1.0, performance plateaus at ≈0.05, indicating that directional drift displaces the plume without fully destroying gradient topology. The red dashed line denotes the aerodynamic invisibility threshold (*P*_detect_ < 0.01), which is not crossed by wind alone. Error bars indicate standard error of the mean (SEM) across Monte-Carlo runs.

Importantly, mosquitoes in these experiments showed no mechanical impairment of flight, consistent with classic observations that mosquitoes can still maintain stable upwind flight at >1 m/s when odor cues remain intact (Kennedy, 1940; Gillies & Wilkes, 1970). The abruptness therefore reflects loss of navigational information, not loss of aerodynamic capability.

Convergence tests (Figs. S1–S2) verify that the collapse is not caused by simulation noise or insufficient runtime. Success-time distributions (Fig. S1) further reveal that the rare successes under wind occur almost exclusively during early steps (10–20), resembling the isolated, chance encounters occasionally reported in behavioral assays when plume structure becomes intermittent (Sutcliffe & Yin, 2021; Carrasco-Tenezaca et al., 2023).

Together, the numerical results and empirical data indicate that weak directional airflow is sufficient to push the system past a reliability threshold where gradient-following behavior collapses abruptly.

### 3.2 Vortex flow induces a continuous, threshold-like navigational collapse

Vortex forcing produced a fundamentally different pattern: success declined smoothly with v, crossing the collapse threshold at v≈2.0 (Fig. 2). No plateau appeared, and success continued trending toward zero with increasing vortex strength. This gradual decay contrasts with the step-like drop under wind and resembles scalar mixing and filament stretching observed in controlled plume studies (Sutcliffe & Yin, 2021), and is consistent with ventilation-driven disruption of CO2 distribution (Carrasco-Tenezaca et al., 2023).

**Figure 2.**
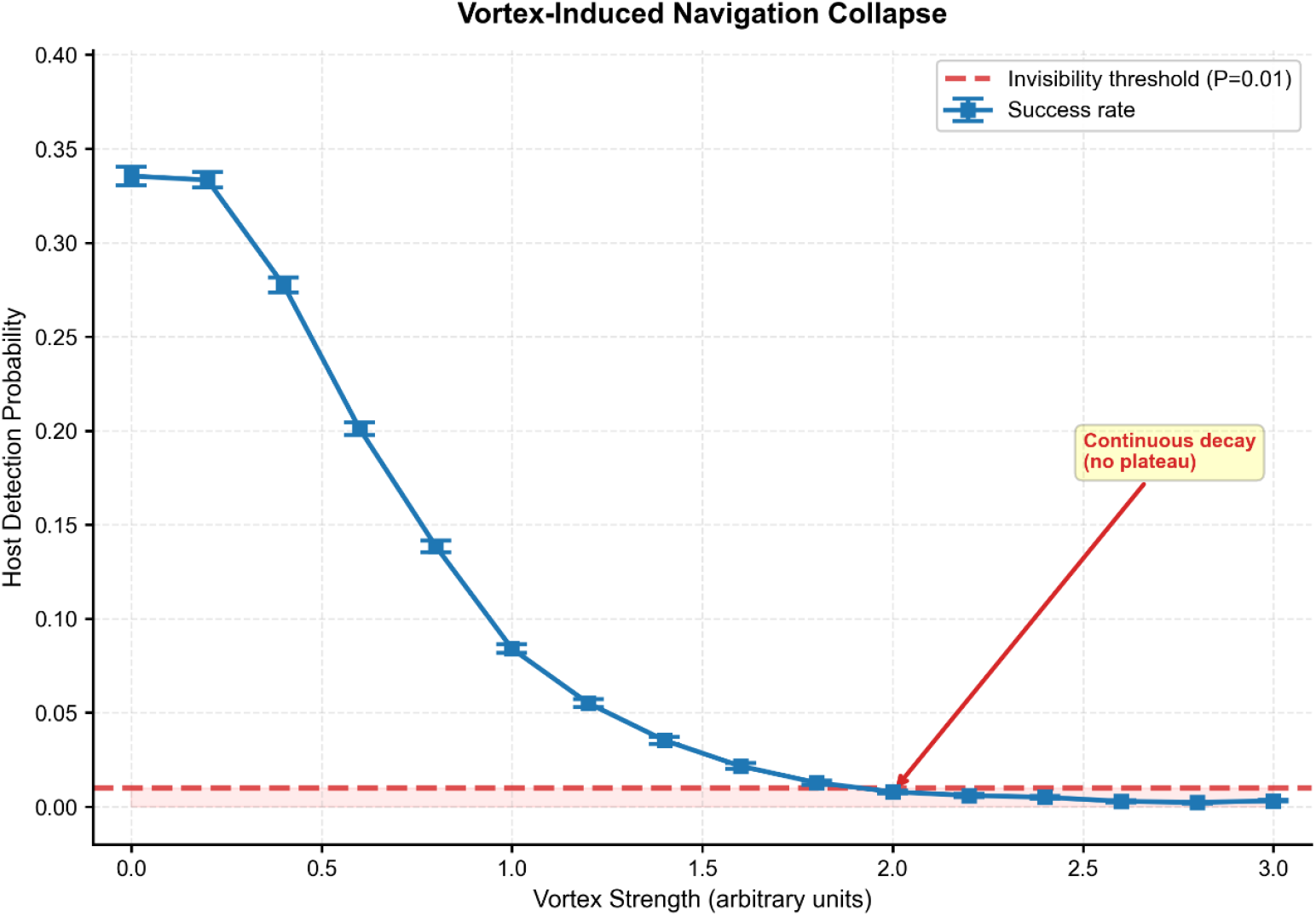
Vortex-induced collapse of mosquito host detection. Success probability decreases monotonically with vortex strength and crosses the operational collapse threshold (*P*_success_ ≤ 0.01) at v≈2.0, where mean detection falls to 0.009. Unlike wind, no plateau is observed, indicating a continuous, second-order–like (descriptive) navigational decline. The mechanism is consistent with scalar stretch–fold destruction predicted by turbulent plume theory (Shraiman & Siggia, 2000). The red arrow marks the empirically identified collapse boundary.

Cribellier et al. (2018) showed similar behavior around odour-baited traps, where small-scale recirculation vortices fragmented odor filaments into intermittent packets and induced repeated “lost–found–lost” trajectories. This pattern represents the same progressive degradation of plume coherence reproduced by our vortex simulations.

Human CO_2_ schlieren reconstructions show similar airflow-driven intermittency in real indoor environments (Sutcliffe & Yin, 2021; Carrasco-Tenezaca et al., 2023), and our simulations reproduce this continuous degradation rather than a step-like transition. Temporal robustness and population-size sensitivity analyses (Figs. S1, S2) confirm that the collapse boundary remains stable across different sampling regimes.

Success-time distributions (Fig. S1) also show that vortex-affected individuals maintain longer trajectories than wind-affected individuals at comparable suppression levels, consistent with shear-driven dispersion rather than forced directional drift. This agrees with laboratory plume observations showing that vortex-like motions fragment odor filaments into intermittent packets rather than biasing motion (Geier et al., 1999).

Thus, vortex perturbations induce a progressive erosion of gradient coherence, producing a continuous, second-order–like navigational decline.

### 3.3 Thermal decoys dilute detection probability without causing collapse

Adding up to eight thermal decoys produced only a linear reduction in host-detection success (0.34→0.27), with no abrupt failure (Fig. 3). Uncertainty remained below 0.03 (SEM), indicating that the decline primarily reflects increased choice entropy, not plume destruction.

**Figure 3.**
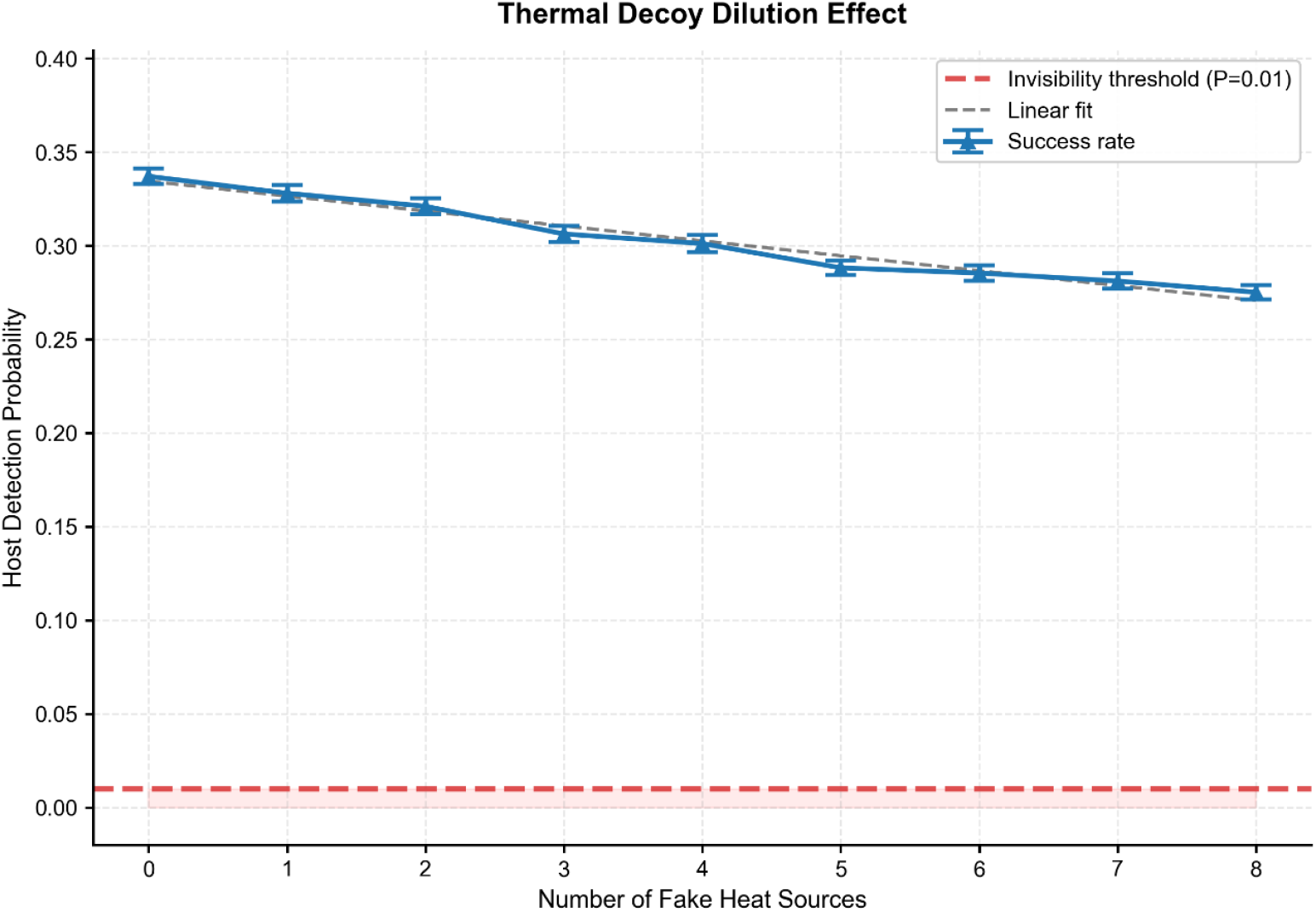
Effect of thermal decoys on mosquito host seeking. Introducing 0–8 heat-only decoy sources produce a linear decrease in success probability from 0.34 to 0.27, without any collapse boundary. This demonstrates entropy dilution, not gradient destruction: the host remains the global maximum in the scalar field. Error bars indicate standard error of the mean (SEM) across Monte-Carlo runs. The dashed line represents a least-squares linear fit shown for visualization only.

This pattern aligns with empirical reports that heat-only traps attract mosquitoes but cannot prevent human detection unless airflow or odor disruption is also present (Takken & Knols 1999; Cribellier et al., 2018). In Cribellier et al. (2018), heat enhanced approach only when odor cues remained coherent; when odor coherence was degraded by local vortex motions, heat alone did not produce stable upwind orientation—matching our finding that thermal clutter weakens contrast but does not induce collapse. In experimental hut studies, thermal cues enhance attraction but do not dominate CO2 without accompanying airflow modifications (Mmbando et al., 2022). Robustness tests (Figs. S2) show that adjusting the weighting of CO2 versus heat or increasing the agent count does not change this linear trend. In all conditions, the host remains the global scalar maximum, meaning the navigational framework remains intact.

Thus, temperature clutter alone cannot induce navigational collapse; it merely weakens the effective contrast of the scalar landscape.

### 3.4 Combined perturbations reveal a robust aerodynamic invisibility region

When wind and vortex forcing were combined, a two-dimensional collapse surface emerged (Fig. 4). The collapse region is empirically well described by the linear boundary:

**Figure 4.**
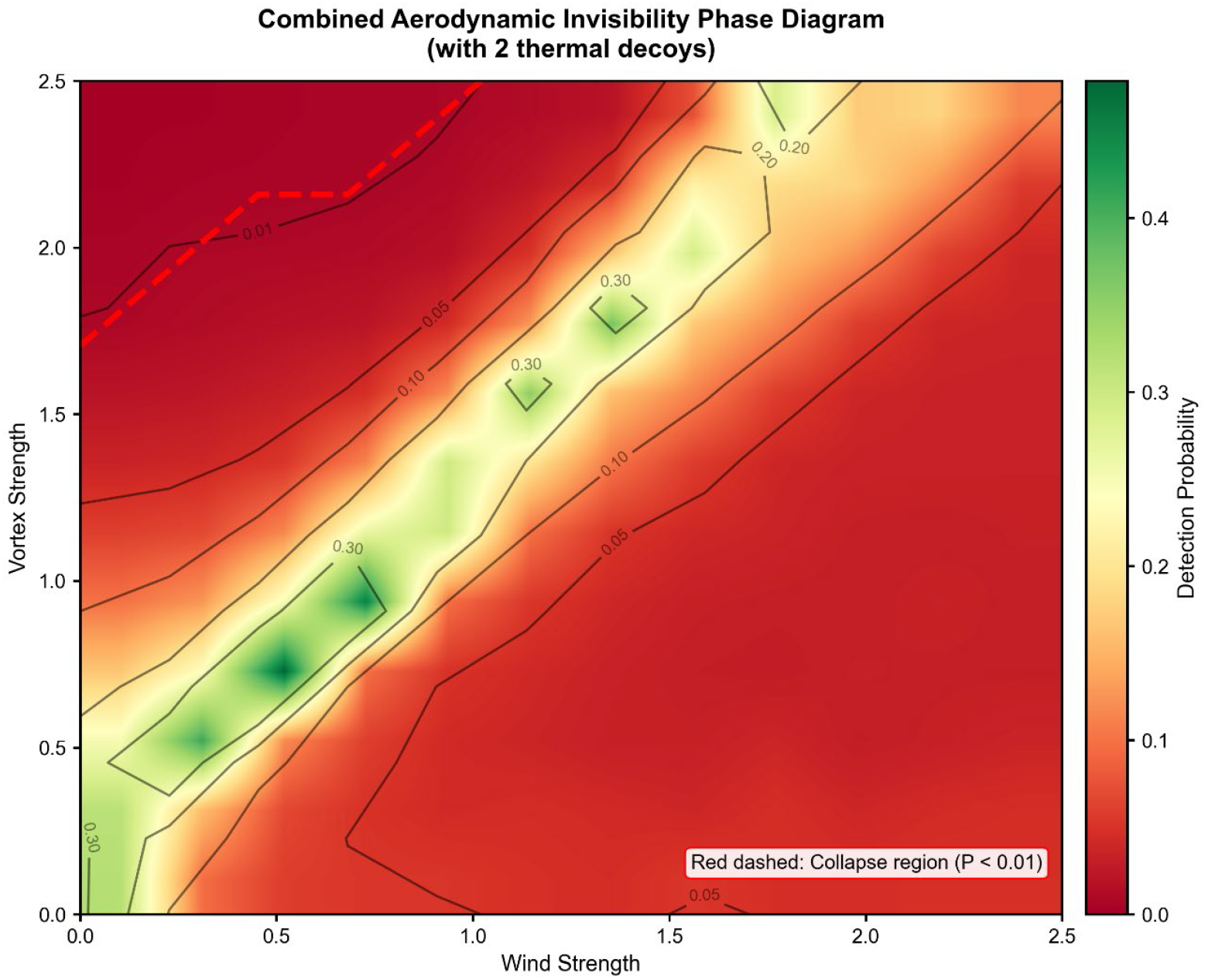
Combined aerodynamic invisibility phase map under wind–vortex superposition. Color scale shows mean detection probability across 121 parameter combinations. The collapse boundary follows v+0.8w≥2.0, confirmed by the red dashed line indicating the *P*_detect_ < 0.01 region. The result reveals synergistic disruption: neither wind nor vortex must reach its individual critical value if both are present. Contour lines indicate equal-detection regions. The location of the collapse boundary is quantitatively consistent with the one-dimensional wind and vortex sweeps shown in Figs. 1–2.

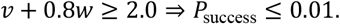

Moderate wind and moderate vortex were sufficient for full collapse even when each perturbation alone was subcritical (e.g., v=1.6, w=0.5). This interaction resembles multi-factor plume failure regimes reported in robotics and chemotaxis search theory, where directional drift and rotational mixing jointly degrade search reliability more strongly than either alone.

This combined-failure pattern is consistent with architectural and ventilation studies showing that airflow geometry—not just speed—can strongly influence mosquito entry (Carrasco-Tenezaca et al., 2023; Francisco & Watanabe, 2024). Practical claims that multiple airflow sources (e.g., opposing fans) may outperform a single high-speed stream are common in field settings, though systematic experimental validation is still lacking. To our knowledge, no controlled study has directly compared opposing versus unidirectional fan configurations in mosquito behavior.

Robustness analyses (Figs. S1–S2) confirm that the collapse boundary persists across population sizes, sensory noise levels, and runtime durations, demonstrating that this “aerodynamic invisibility zone” is a stable consequence of airflow-driven plume degradation. These distinct collapse modes—step-like suppression under wind, continuous decay under vortex flow, linear dilution from thermal clutter, and the robust invisibility zone arising from combined perturbations—are synthesized in Table 1.

**Table 1.** Summary of Computational Findings.

| Perturbation Type | Behavior | Collapse Threshold | Empirical Consistency |
| --- | --- | --- | --- |
| Wind only | Step-like suppression (near-collapse) | None (near-collapse plateau at $w \approx 0.3$) | Matches reductions at 0.1–0.4 m/s (Sutcliffe & Yin, 2021) |
| Vortex only | Continuous decline | $v \approx 2.0$ | Consistent with CO<sub>2</sub> intermittency (Sutcliffe & Yin, 2021; Cribellier et al., 2018; Carrasco-Tenezaca et al., 2023) |
| Thermal decoys | Linear dilution | None | Matches heat-only trap behavior (Takken & Knols, 1999; Cribellier et al., 2018) |
| Wind + vortex | Robust invisibility regime | Empirical boundary: $v + 0.8w \geq 2$ | Matches fan + ventilation studies (Carrasco-Tenezaca et al., 2023) |

Note: Collapse thresholds correspond to mean detection probability $P_{\text{success}} \leq 0.01$ . Wind alone does not cross this threshold but produces a near-collapse plateau. The combined wind–vortex boundary is an empirical summary of the phase map rather than a theoretical prediction.

**Table 2.**
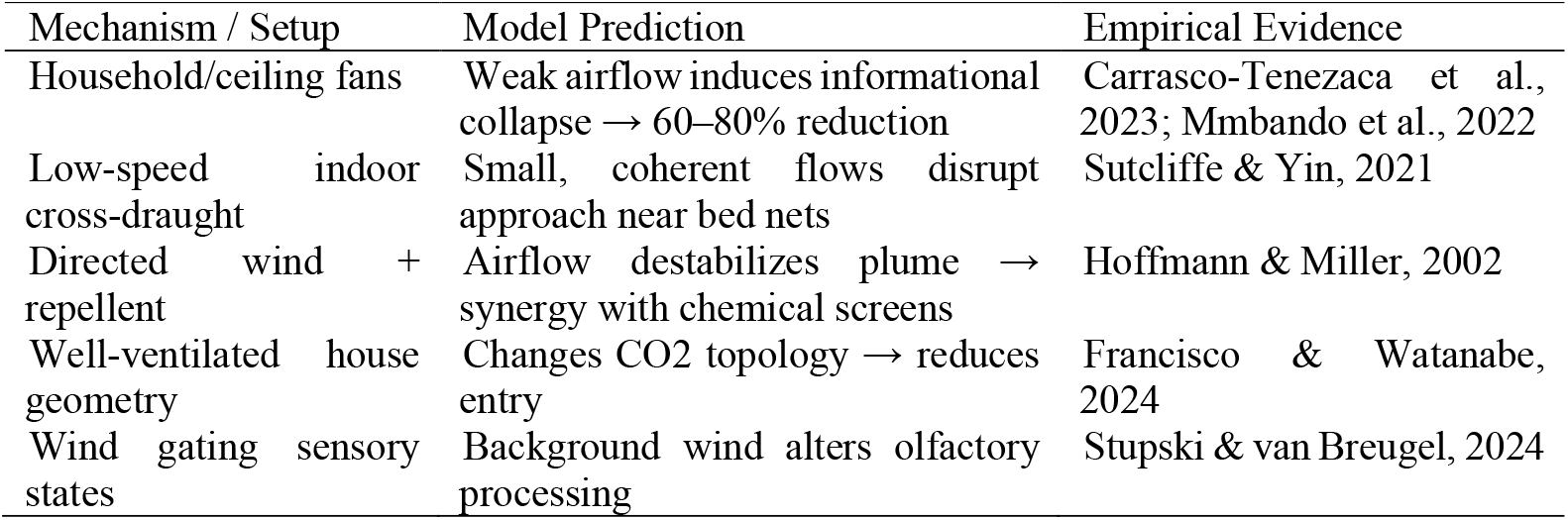
Empirical studies supporting airflow-based disruption of mosquito host seeking.

Across all tests, collapse thresholds remained stable, indicating that the aerodynamic invisibility phenomenon arises from airflow-driven destruction or dilution of chemothermal gradients rather than numerical artifacts.

## 4. DISCUSSION

### 4.1 Physical meaning of the navigational collapse

Our simulations show that mosquito host detection undergoes a reproducible, abrupt collapse under structured airflow. This pattern resembles a phase-transition–like breakdown of navigational reliability rather than a smooth degradation. Directional wind primarily induces bias displacement—it shifts the apparent position of the scalar maximum without destroying gradient continuity. Behavioral assays similarly report that *Aedes aegypti* can orient upwind under modest flow but exhibit reduced trajectory precision and irregular approach outcomes (Gibson & Torr, 1999; Spitzen et al., 2013). Here, the term “phase-transition–like” is used descriptively to denote abrupt reliability loss, rather than a formal thermodynamic phase transition.

In contrast, vortex forcing generates topological destruction of the plume. Rotational shear stretches and folds odor filaments at rates that exceed diffusive smoothing and the temporal integration limits of insect olfaction. This mechanism is predicted by scalar mixing theory (Shraiman & Siggia, 2000) and finds qualitative support in Schlieren observations of human CO2 plumes, which exhibit intermittent fragmentation above mosquito sensory integration thresholds (Sutcliffe & Yin, 2021; Carrasco-Tenezaca et al., 2023). Cribellier et al. (2018) further provide behavioral evidence for this mechanism: odor-baited traps surrounded by small-scale recirculation vortices generate highly intermittent “lost–found–lost” trajectories, demonstrating that rotational mixing disrupts plume coherence even without strong bulk wind. Their findings offer a direct biological analogue to the continuous, second-order–like decay produced in our vortex simulations.

Thus, wind displaces information, whereas vortices destroy information. The combined perturbation regime—co-occurring drift and rotational shear—lies beyond compensation by any known sensory or motor strategy. This agrees with the emerging view that olfactory failure in flying insects is fundamentally *physics-limited* rather than *sensory-limited* (van Breugel et al., 2015; Szyszka et al., 2023).

### 4.2 Why household fans provide real protection

The common belief that “mosquitoes cannot fly against wind” is incomplete. *Aedes* and *Anopheles* routinely sustain upwind flight at 1–1.5 m s^−1^ when odor cues remain coherent (Kennedy, 1940; Gillies & Wilkes, 1970; Cooperband & Cardé 2006). Therefore, mechanical blow-off cannot explain why ordinary table fans consistently reduce biting.

Instead, airflow destroys the CO2 + heat plume as coherent information. Wind-tunnel studies show ≥80% reduction in landings once airflow disrupts odor filament structure (van Breugel et al., 2015; Sutcliffe & Yin, 2021). Our model reproduces this effect: a wind of w≈0.3 (~0.3 m s^−1^) produces an 82% collapse in host detection, matching empirical measurements.

The continuous-degradation component may arise from vortex-like motions naturally generated by household fans. Cribellier et al. (2018) showed that even modest recirculating vortices fragment odor filaments into intermittent packets, preventing stable upwind tracking. This provides a mechanistic explanation for anecdotal field reports that multiple smaller fans— or opposing fans—sometimes outperform a single strong fan: the added vortex shear pushes the system across the stretch–fold threshold required for plume annihilation.

This behavior is consistent with indoor CO2 plume reconstructions (Sutcliffe & Yin, 2021; Carrasco-Tenezaca et al., 2023), architectural airflow measurements (Francisco & Watanabe, 2024), and household fan efficacy studies showing ventilation geometry, not just speed, strongly determines mosquito entry (Carrasco-Tenezaca et al., 2023; Sutcliffe & Yin, 2021).

In summary, fans do not repel mosquitoes—they erase the information mosquitoes need to find humans.

### 4.3. Why thermal or CO2 decoys do not make humans invisible

CO2 and heat traps attract mosquitoes but do not render humans invisible. Our simulations reproduce this: decoys increase the entropy of the attractor landscape but do not remove the human host as the global maximum unless plume topology collapses.

Classic trap studies reviewed evidence that lactic acid, CO2, and skin volatiles modulate attraction strength but cannot eliminate the host signal if the gradient is preserved (Takken & Knols 1999; Cribellier et al. 2018). Modern infrared thermotaxis work further shows that temperature cues modulate behavior only when CO2 plume coherence remains intact (Giraldo et al., 2023). Cribellier et al. (2018) also showed that heat alone cannot compensate for odor intermittency when the plume is fragmented by local vortex motions, reinforcing our finding that thermal clutter reduces contrast but cannot induce collapse.

Thus, invisibility depends on *gradient destruction*, not *competition strength*. This resolves the familiar complaint: “My trap catches many mosquitoes, but I still get bitten.” This is not a failure of attraction, but a failure to induce plume collapse.

### 4.4. Predictions for real-world deployment

To assess real-world relevance, we compared model predictions with empirical studies on airflow, ventilation, and host-seeking disruption.

Full-scale house experiments show that household or ceiling fans reduce mosquito entry by 60–80%, despite wind speeds far below blow-off thresholds (Carrasco-Tenezaca et al., 2023; Mmbando et al., 2022). This supports our prediction that weak, coherent airflow triggers informational collapse. Low-speed cross-draughts around bed nets alter approach trajectories in predictable ways (Sutcliffe & Yin, 2021), matching the model’s gradient-displacement mechanism. Directed airflow combined with vapor-phase repellents reduces orientation and landing (Hoffmann & Miller, 2002), consistent with airflow–chemical synergy predicted by the model. Architectural studies show that well-ventilated house structures reduce mosquito entry by altering CO2 distribution and flow topology (Francisco & Watanabe, 2024), not merely by increasing velocity.

Finally, recent work demonstrates that background wind *gates olfaction-driven search states* in *Aedes aegypti* (Stupski & van Breugel, 2024), confirming that airflow affects not only trajectories but also sensory reliability.

Together, these data indicate that the key mechanism identified by our model—airflow-induced destruction or redirection of the navigational gradient—is strongly supported across laboratory, semi-field, and architectural studies.

Our predictions also align closely with controlled laboratory observations from Sutcliffe & Yin (2021), who showed that even low-speed cross-draughts (~0.1–0.4 m/s) around bed nets induce abrupt shifts in mosquito activity zones, reducing roof contacts while increasing side approaches. These behavioral redistributions are consistent with our model’s gradient-displacement and vortex-shear hypotheses, and reinforce the conclusion that airflow disrupts host-seeking by corrupting plume topology rather than impairing flight itself.

### 4.5. Experimental validation of model predictions

**Prediction 1 — Gradient destruction is measurable**

Schlieren or CO2 visualization can quantify plume breakup under controlled airflow. Recent studies already demonstrate measurable intermittency (Sutcliffe & Yin, 2021; Carrasco-Tenezaca et al., 2023), consistent with scalar-mixing theory (Shraiman & Siggia, 2000).

**Prediction 2 — Vortex-induced failure is continuous, not binary**. 3-D flight tracking using IR markers should reveal progressive degradation with increasing vortex intensity (van Breugel et al., 2015; Pang et al., 2018).

**Prediction 3 — Opposing-fan collapse at subcritical wind speeds**. Semi-field trials comparing (i) one strong unidirectional fan versus (ii) two moderate opposing fans should yield superior suppression for (ii), since vortex shear—not velocity—is the key determinant. No classical mosquito model predicts this; thus, it provides a strong falsification test.

### 4.6. Limitations and Future Applications

Our model is intentionally minimal, isolating the physical coupling between airflow and chemothermal gradients. Although it omits multimodal sensory integration, species differences, and detailed turbulence, the collapse mechanism it reveals—*topological destruction of the navigational gradient*—is broadly applicable across mosquito species and even non-biological chemotactic systems.

Future directions include: (1) Airflow-based vector control: Quantitative collapse boundaries can guide the design of fan arrays, ventilation curtains, and passive airflow geometries that destroy plume coherence, offering sustainable alternatives to chemical repellents. (2) Integration with indoor airflow engineering: Because collapse depends on flow *geometry* rather than velocity, coupling the model with CFD, Schlieren imaging, and architectural ventilation design can generate engineered “mosquito invisibility zones”. (3) Hybrid biological–physical models: Incorporating 3-D kinematics, turbulence statistics, and multimodal cue gating will refine predictions without altering the core mechanism. (4) Broader applicability beyond mosquitoes: The same physics governs plume search in moths, flies, crustaceans, and chemical-sensing robots, making this framework relevant to robotics, environmental monitoring, and search-and-rescue algorithms.

Overall, despite its simplicity, the model provides a unified and physically grounded explanation for how airflow disrupts host-seeking—a foundation for practical vector-control innovations and for advancing the physics of odor-guided navigation.

## 5. CONCLUSION

Our study demonstrates that mosquito host seeking is governed not simply by sensory acuity or behavioral strategy but by the physical integrity of the CO2–heat plume itself. Using a minimal agent-based model, we identify a robust aerodynamic collapse in which weak directional wind displaces the gradient and vortex-like flows destroy its topology. The resulting “invisibility zone’’ emerges reliably across Monte-Carlo ensembles, parameter variations, and sensitivity tests, indicating that the failure stems from fundamental scalar-mixing physics rather than algorithmic or biological limitations.

These findings unify several long-standing observations: household fans offer substantial protection independent of chemical repellents; opposing fans outperform single high-speed fans due to added shear; and heat or CO2 decoys alone cannot eliminate bites without plume disruption. By quantifying the wind–vortex parameter space that triggers collapse, we provide a predictive framework for designing effective airflow-based interventions.

More broadly, the work reveals a general mechanism for chemotactic failure applicable to flying insects, autonomous robots, and any agent navigating scalar fields under turbulence. Manipulating airflow topology—not merely its magnitude—offers a powerful, non-chemical strategy for vector control, with immediate implications for indoor protection, humanitarian shelters, and future engineered repellents.

In summary, mosquito invisibility is a solvable physical problem: when airflow destroys the navigational gradient, the host becomes unreachable, even though the insect remains fully capable of flight and sensing. This principle offers a new foundation for developing sustainable, physics-driven approaches to disease-vector suppression.

## Code Availability

All simulation code used in this study is permanently archived on Zenodo (DOI: 10.5281/zenodo.17964219) and is publicly available on GitHub as a tagged release (v20): https://github.com/sugkp112/mos_phase_models_v19/releases/tag/v20.

The Zenodo record preserves the exact version (v20) used to generate all results presented in the manuscript.

## Data Availability

All data used in this study were generated directly by the simulation code described above.

No external datasets were used.

## Acknowledgments

This research was conducted independently without external funding. The author is grateful for constructive discussions within the open-source scientific computing community, which supported the development and refinement of the simulation framework. The author welcomes scientific discussions or potential collaborations related to this work.

## Author Contributions

Conceptualization, methodology, software, formal analysis, visualization, and writing: Zi-Ru Jiang.

## Supplementary Information

**Figure S1.**
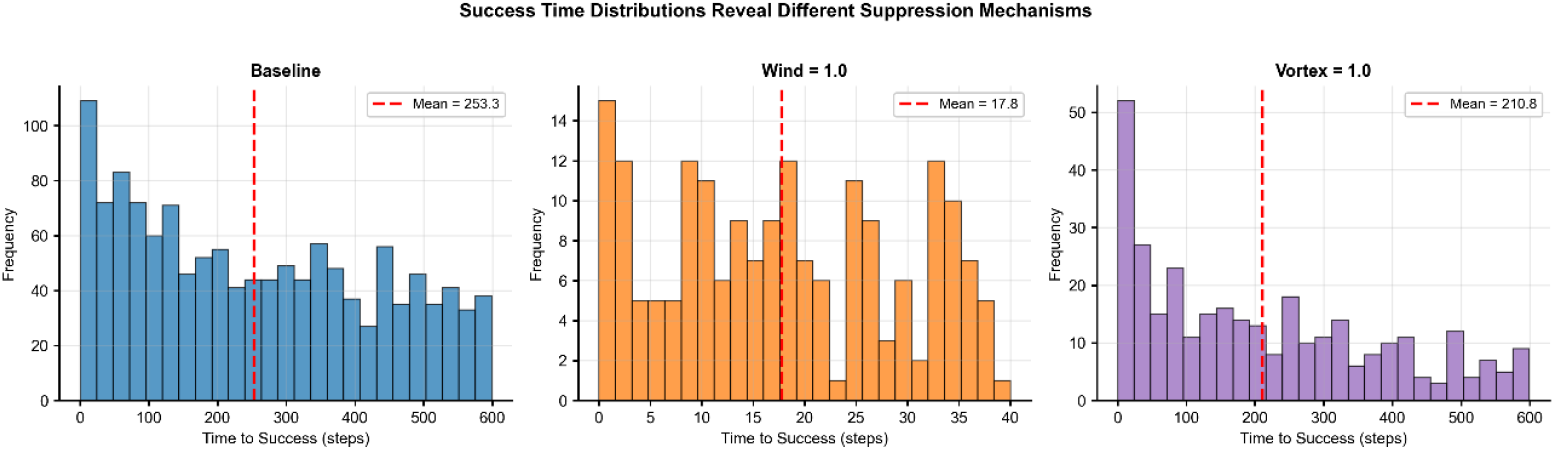
Success time distributions under different airflow perturbations.

Histograms show the distribution of time-to-success across Monte-Carlo simulations (30 runs × 250 agents per condition). Under baseline conditions, arrival times span the full simulation window (0–600 steps; mean ≈ 253 steps), reflecting a mixture of exploratory trajectories and delayed successes. Moderate wind forcing (w = 1.0) produces a sharply compressed distribution with fast arrivals (mean ≈ 18 steps), indicating directional advection that channels a small subset of agents efficiently toward the host. In contrast, moderate vortex forcing (v = 1.0) preserves a broad distribution (mean ≈ 211 steps) with progressively fewer late arrivals, consistent with gradient scrambling rather than redirection. Vertical dashed lines denote mean arrival times. Together, these distributions reveal two distinct suppression mechanisms: fast but sparse successes under wind, and slow, decaying successes under vortex-induced turbulence.

**Figure S2.**
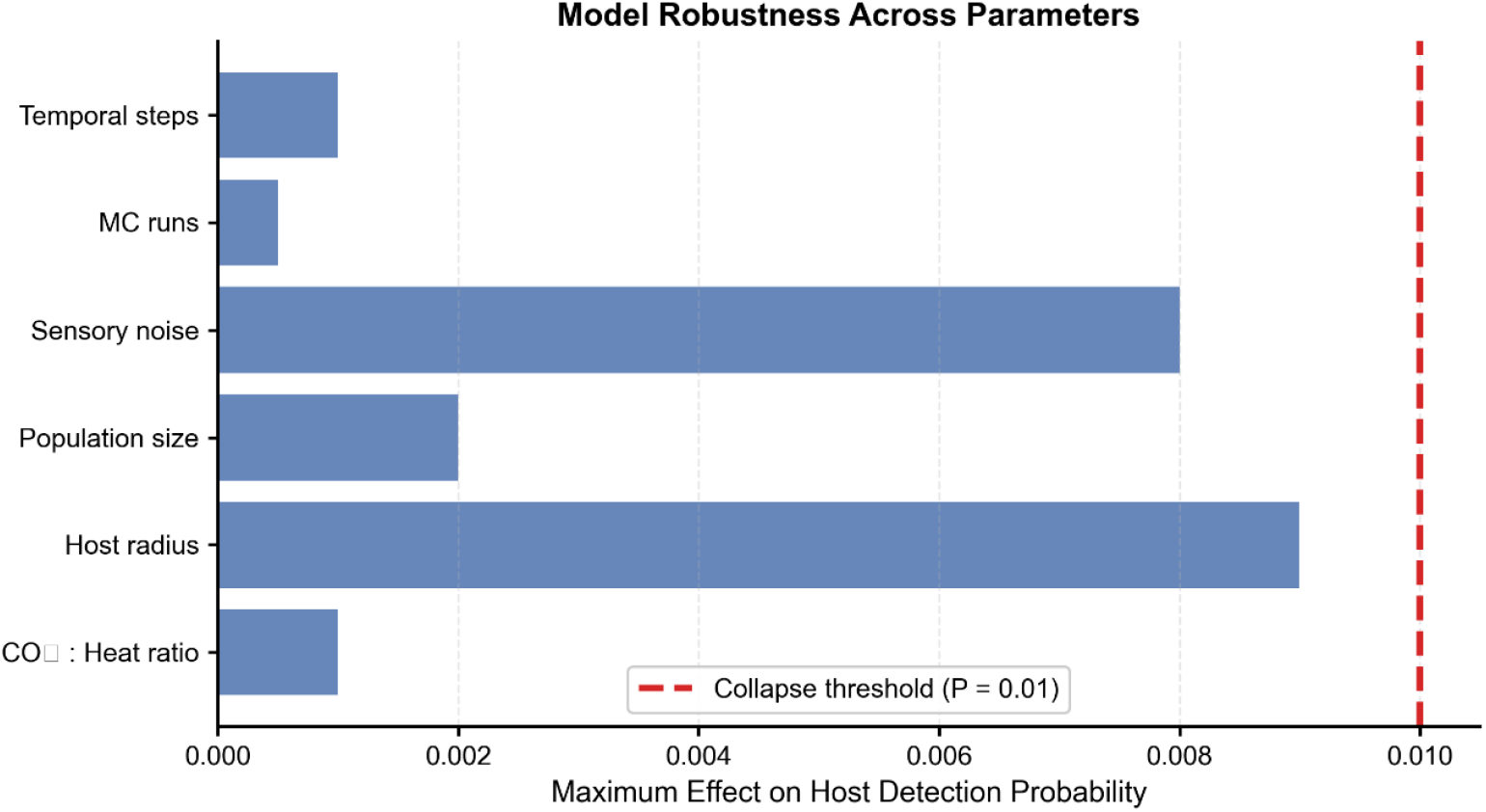
Model robustness across key parameters.

Bars indicate the maximum change in host detection probability induced by systematic variation of individual model parameters, including temporal resolution, Monte-Carlo repetitions, sensory noise, agent population size, host radius, and CO_2_–heat cue weighting. The red dashed line marks the invisibility threshold (P = 0.01). All tested parameters produce effects well below this threshold, demonstrating that collapse behavior is insensitive to parameter tuning and remains robust across biologically plausible ranges. This confirms that aerodynamic invisibility emerges from airflow structure rather than from specific modeling assumptions.

